# Category-Selective Visual Regions Have Distinctive Signatures of Connectivity in Neonates

**DOI:** 10.1101/675421

**Authors:** Laura Cabral, Leire Zubiaurre, Conor Wild, Annika Linke, Rhodri Cusack

## Abstract

The development of the ventral visual stream is shaped both by an innate proto-organization and by experience. The fusiform face area (FFA), for example, has stronger connectivity to early visual regions representing the fovea and lower spatial frequencies. In adults, category-selective regions in the ventral stream (e.g. the FFA) also have distinct signatures of connectivity to widely distributed brain regions, which are thought to encode rich cross-modal, motoric, and affective associations (e.g., tool regions to the motor cortex). It is unclear whether this long-range connectivity is also innate, or if it develops with experience. We used MRI diffusion-weighted imaging with tractography to characterize the connectivity of face, place, and tool category-selective regions in neonates (N=445), 1-9 month old infants (N=11), and adults (N=14). Using a set of linear-discriminant classifiers, category-selective connectivity was found to be both innate and shaped by experience. Connectivity for faces was the most developed, with no evidence of significant change in the time period studied. Place and tool networks were present at birth but also demonstrated evidence of development with experience, with tool connectivity developing over a more protracted period (9 months). Taken together, the results support an extended proto-organizon to include long-range connectivity that could provide additional constraints on experience dependent development.

## Introduction

The development of the ventral visual stream depends upon both its innate architecture and on visual experience. The role of innate architecture is apparent in the stereotypical locations of category-selective regions ^1–5^. But, experience is also essential for typical development ^6,7^, and category-selective regions can develop even for modern socio-cultural artifacts like writing^8–10^.

Current evidence suggests an innate proto-organization within the ventral visual stream that shapes experience-dependent development. The fusiform face area (FFA) ^4^, for example, receives stronger visual input from the fovea and lower spatial frequencies ^11^, leading to the acquisition of representations that are particularly suited for processing faces. Recently, it was found that this proto-organization is present at 6-27 days, as a face-selective region had stronger functional connectivity with foveal V1 than peripheral V1; while the reverse was seen for a scene region ^12^.

While there is evidence early in life for proto-organization within the ventral visual stream, less is known about the connections from the ventral stream to other areas of the brain. By adulthood, different category-selective regions have distinct connectivity patterns ^13,14^. Such long-range connections are thought to encode the cross-modal, motoric and affective associations characteristic of rich semantic categories ^15–17^. As concrete examples, seeing a silent video of a dog barking evokes the representation of its sound in auditory cortex ^18,19^, and for tools and objects, category representations in the ventral stream are integrated with action representations in the dorsal stream ^20–22^. The importance of long-range connections in semantics was recently demonstrated by multivariate decoding of white matter pathways in brain-injured patients with semantic deficits ^23^.

At present, it is unclear whether this long-range connectivity develops with experience or is largely innate. It could be that it reflects experience, through the statistical learning of associations. According to this model, for example, in the first stage of learning face selectivity might mature in a bottom-up way through biased input ^6,11,12,24^. The auditory system might be maturing from its own proto-organization, perhaps due to selectivity to temporal modulations, to develop voice-selective regions ^25^. Subsequently, as faces co-occur with voices, there might be Hebbian strengthening of the pathway between face and voice selective regions. An alternative model is that the distinct long-range category-selective connectivity is innate. According to this model, a connection that exists at birth between the eventual face- and voice-selective regions could shape the development of their selectivity.

To differentiate these two models, we used neuroimaging to investigate the maturity of long-range structural connectivity of category-selectivity regions in neonates to 9-months-old infants. To measure connectivity, we used diffusion-weighted imaging and tractography. We extracted the characteristic signatures of connectivity of three category-selective regions in adults using a machine learning approach, and then tested for generalization to infants in an in-house dataset. To address limitations, we then extended our analysis using neonatal data from the developing human connectome project.

## Results

### Experiment 1

#### Category-Selective Regions Have Distinctive Signatures of Connectivity in Young Infants

In order to measure connectivity, diffusion-weighted images were obtained from 14 adults and 11 infants. The images from each individual were warped to a standard template space using the fractional anisotropy maps. Even in the youngest infant the major white matter structures are apparent in these maps providing substantial information to guide registration. Example slices from the infants are shown in **Figure 1**.

**Figure 1.**
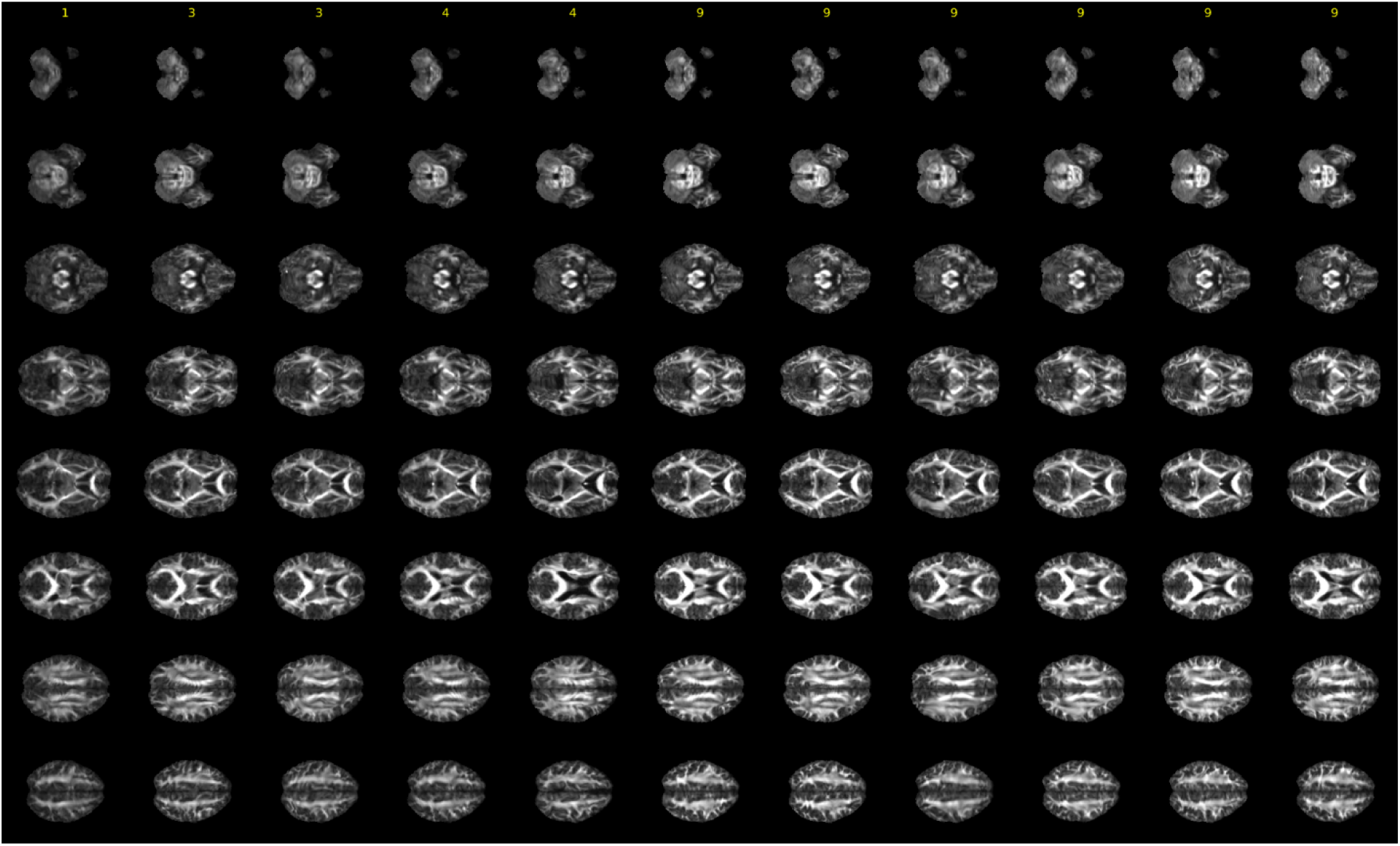
Diffusion weighted images (FA) for each of the 11 infants. Each column represents an individual infant and each row represents a volume. Across ages, there is good registration across the sample, as illustrated by the similarity of each volume among infants.

Our goal was to characterise the brain-wide pattern of connectivity of each category selective region. To do this, probabilistic tractography was performed using seed and target regions taken from the parcellation by the Human Connectome Project (HCP) ^26^. Each voxel in the ventral visual stream region of interest defined by an HCP template^26^ was used as a seed, while the brain areas outside the ventral stream acted as targets for tractography (see **Figure 2**). To define category-selective regions in the ventral stream, contrast maps from the HCP fMRI localizers were used to determine the regions within the parcellation that were most selective for face, places and tools^26^. These regions were the fusiform complex, the ventromedial visual area 2, and ventromedial visual area 3, respectively. Regions of interest are displayed in **Figure 2a.**

**Figure 2.**
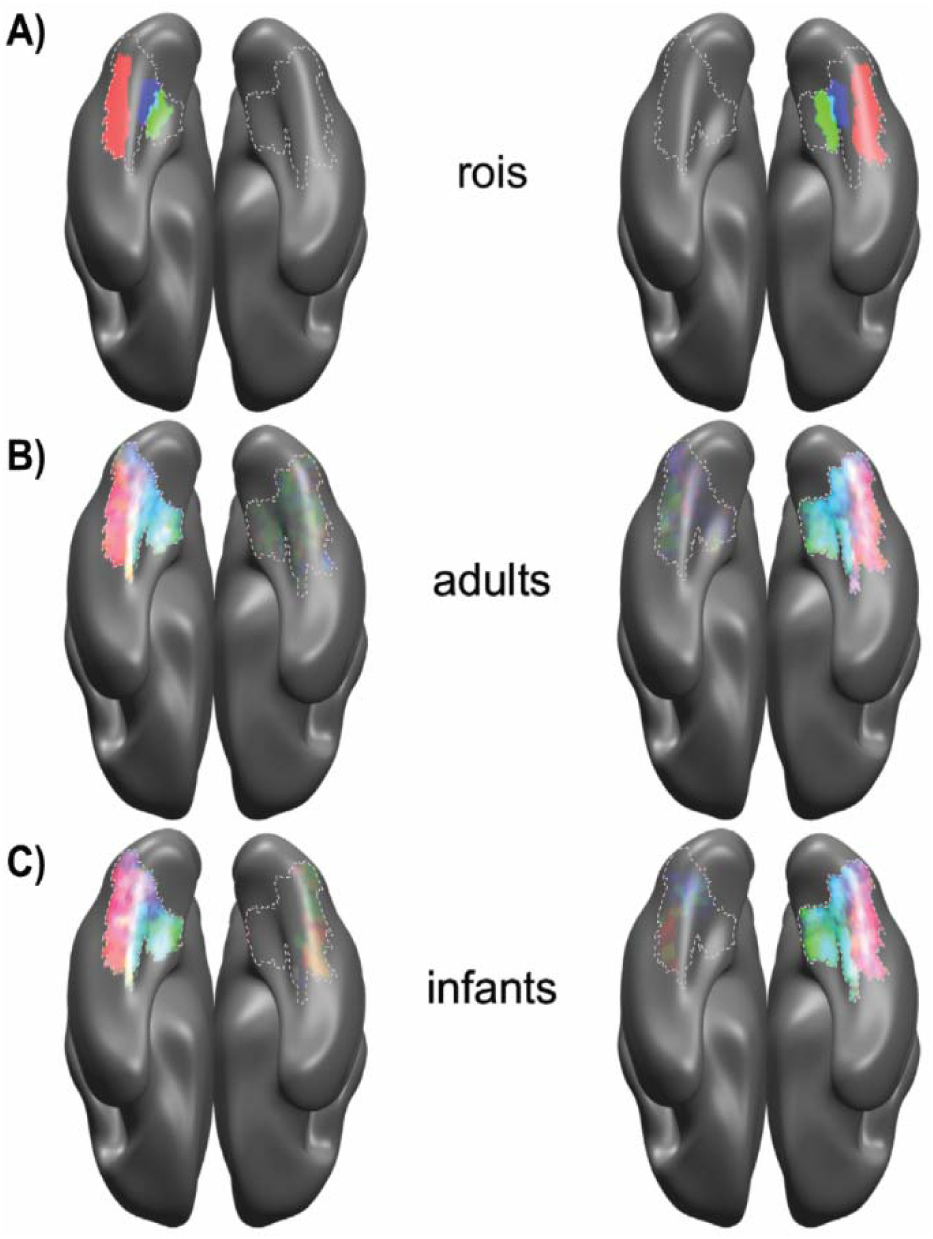
This figure shows the regions used for classification and the voxels selected by the classifiers as part of the face, place, and tool regions of interest based on their structural connectivity with other brain regions. A) Regions from the HCP parcellation that were most selective for faces, tools and places (red, blue and green, respectively) in the left and right hemispheres (left and right columns). Dotted outlines represent the ventral stream seed region, as defined by the HCP (see methods) B) Voxels identified by a linear-discriminant classifier as selective for faces, places, and tools in adult participants (N=14), based on their distinctive signature of structural connectivity with the rest of the brain. Classification was performed separately for the left and the right hemisphere, using leave-one-subject-out cross validation. Group average overlay maps are shown with the same color mapping as (A). C) The distinctive signatures of structural connectivity were also present in infants (N=11), as shown by voxels identified as category-selective by a linear-discriminant classifier trained on adult connectivity and tested in infants.

The connectivity pattern for the category-selective regions were then probed using six linear-discriminant classifiers, one for each visual category in each hemisphere. Using leave-one-subject-out cross-validation, a classifier was trained to differentiate voxels from the category selective regions from the other voxels in the ventral stream, based on their structural connectivity with the rest of the brain. The classifier’s performance was then tested on the left-out subject. Using signal detection theory, d-primes were calculated for each participant, to determine how sensitive the classifiers were in locating voxels in the face, place and tool regions. All three regions could be robustly localized in adults (*t*(13)=26.26, *p*<0.001, t(13)=22.35, *p*<0.001, t(13)=17.17, *p*<0.001) **(Figure 2b**). Classification performance is quantified in **Figure 3a**, which shows the d-primes for classification of the imaging data.

**Figure 3.**
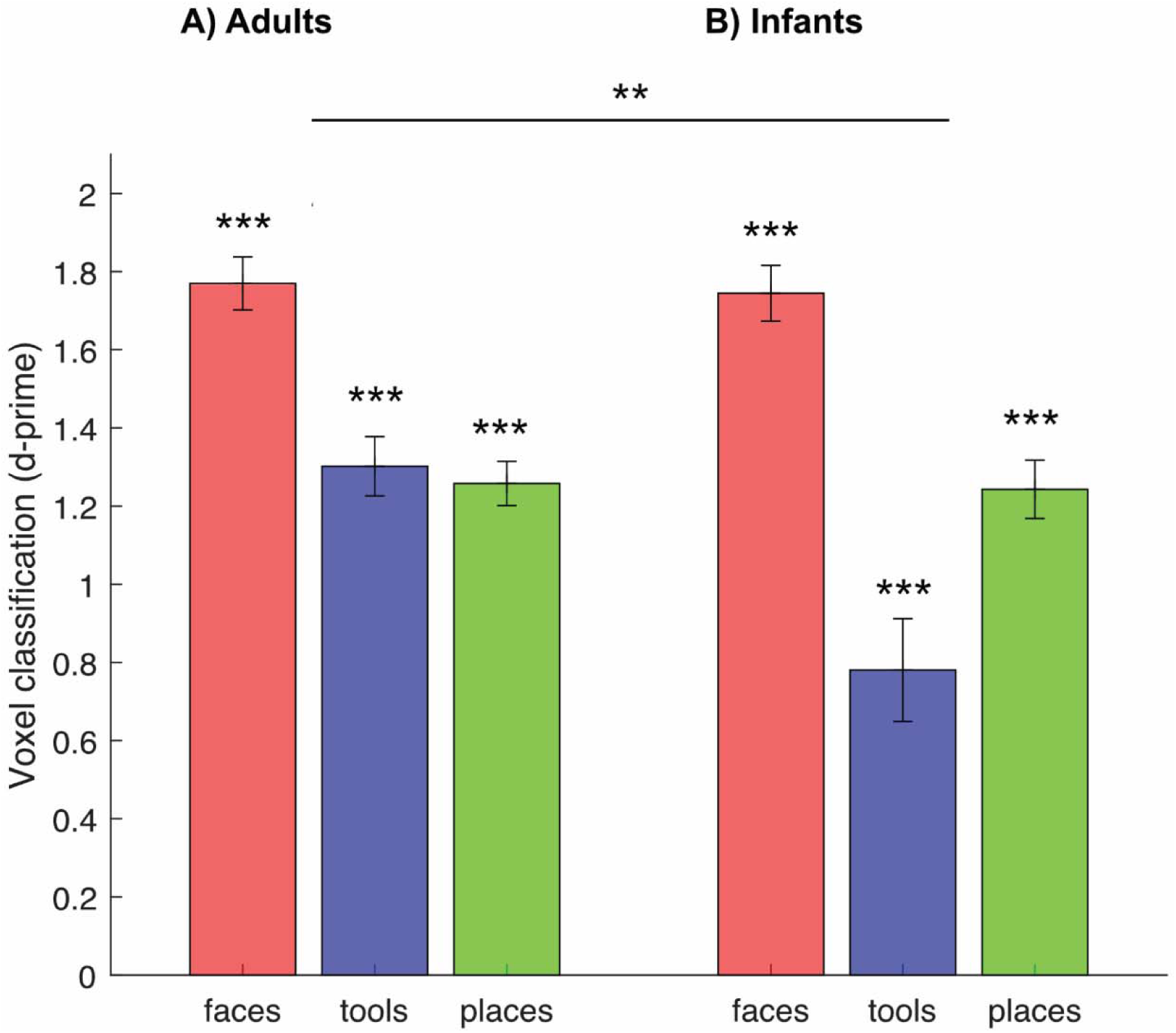
a) Voxel classification performance for the adults (N=14) quantified using d-prime, collapsed across hemispheres. b) Voxel classification performance for the infants (N=11) again measured using d-prime. All regions were robustly localized in infants and adults but there was a significant difference in detection accuracy between the infant and adult tool region, demonstrating the immaturity of connectivity for the tool region during infancy. The mean +/- one standard error across subjects is shown.

In order to characterize the connectivity of the infant ventral stream, probabilistic tractography was also performed on the infant diffusion data, using the same seed and target regions as in adults (see Methods for details of two-stage adult-to-infant normalization procedure). To determine whether the category-selective regions that were present in adults were present in infants, linear discriminant classifiers were trained on the entire adult dataset in the manner described above. These classifiers were then tested on the infant data. The classifiers localized all three regions in the infants (t(10)=24.47, *p*<0.001, *t*(10)=16.54, *p*<0.001, *t*(10)=5.95, p<0.001) **(Figure 2c)**. However, there was a category-by-group interaction (*F*(2,46)=6.64, *p*<0.01). Post-hoc tests showed this was because the face and place regions were as strongly detected in infants as they were in adults (*t*(23)=0.165, *p>0.05, t*(23)=0.257, *p>0.05*), but the tool region was detected with greater accuracy in adults than in infants *(t*(23)=3.62, *p*<0.01**) (Figure 3b)**. Finally, in order to examine the developmental trajectory of the networks, infant age and classification accuracy (d-primes) were correlated--only the tool network underwent significant change over the first 9 months of postnatal life (faces: *r(*9)=-0.03, *p>0.05*; tools: *r*(9)=0.75, *p*<0.01; places: *r*(9)=-0.01, *p>0.05*) **(Figure 4)**.

**Figure 4.**
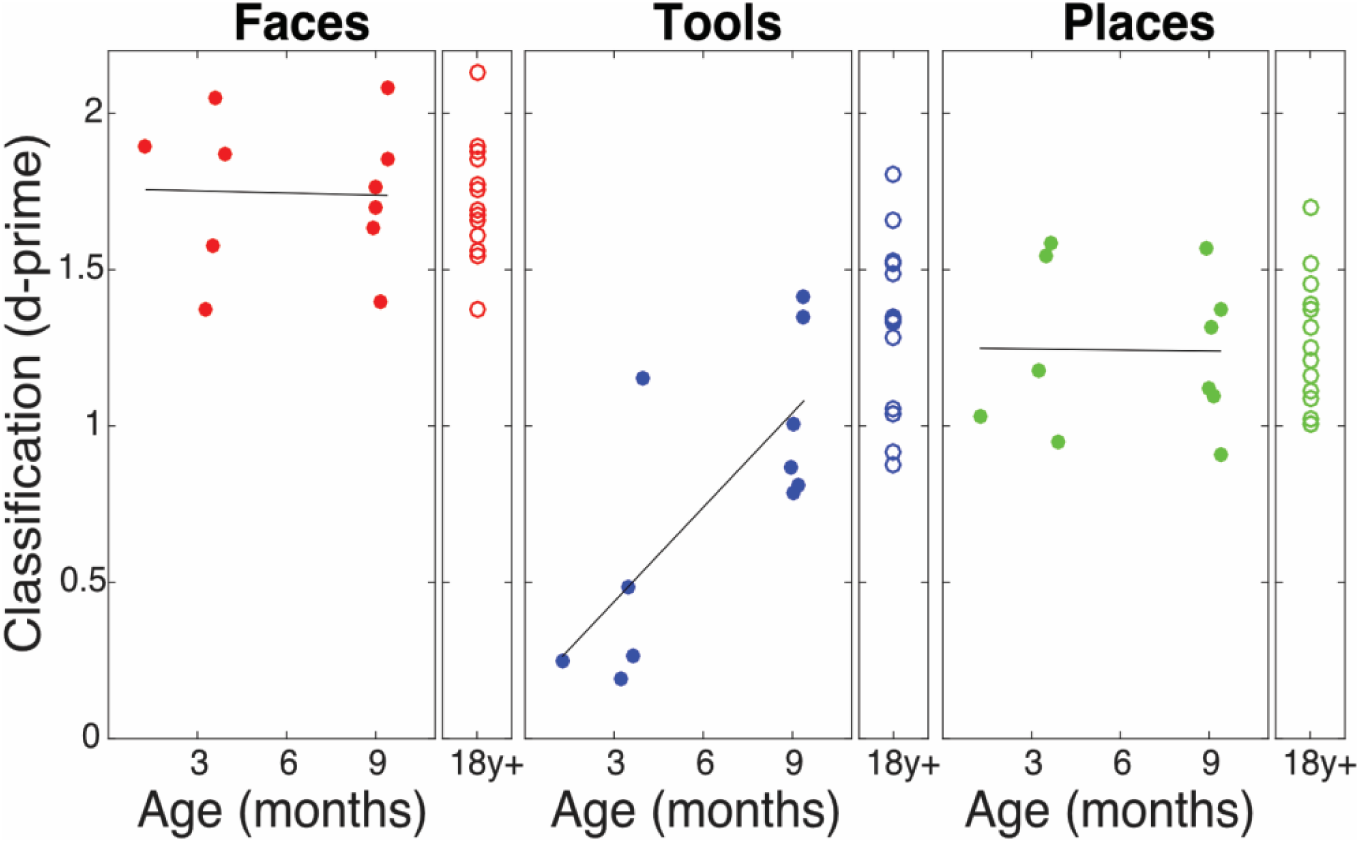
The relationship between the age of participants (14 adults and 11 infants) and classification accuracy (d-prime) with best-fit lines. Only tool classification had a significant relationship with age, demonstrating the maturation of the distinctive connectivity of the tool network over the first year of postnatal life.

These results suggest the connectivity of the tool region develops later than that of the face and place regions, but we also examined an alternative explanation. Could it be that tractography is more difficult in infants than adults, because of their lower signal-to-noise or smaller brains, and that identification of voxels in the tool region is more sensitive to this? Two analyses were conducted to investigate this. First, in adults, the detection of the tool-selective voxels was no worse than detection of the place-selective voxels and performance was not at ceiling (Fig 2A), suggesting that detection of the tool region is not intrinsically more difficult. Second, we compared region size, which may affect performance more strongly in smaller infant brains; the place and tool regions were the same size in one hemisphere and were less than 10 voxels different in the other. The hit rate for the place and tool region was also not significantly different in adults (*t*(13)=-1.04, *N.S*).

We next examined which target regions of connectivity most strongly influenced each of the classifiers in adults (see Supplementary Table 1). We chose to look at the adult data because the classifiers were trained on it, thus defining the regions with high weights that were most important for linear classification. For the place area, this was connectivity to a network strongly associated with navigation, including the hippocampus, parahippocampal areas, and the entorhinal cortex ^2,27^. The face region’s strongest-weighted structural connections were to area PH, which is strongly deactivated in the HCP functional face contrasts ^28^. Additional strong connections in the face network include area PGp and area PGS which are a part of the inferior parietal lobe. It is possible that these are connected to the face region by the inferior longitudinal fasciculus ^29^; this connectivity pattern is substantiated with strong functional connectivity from area PGp to other high-order visual areas ^26^. Finally, although the occipital face area (OFA) and FFA have previously shown strong connectivity, the Glasser et al. (2016) parcellation does not have a specific OFA region. The OFA is classified as part of the PIT Complex, classified as part of the ventral visual stream, so this approach did not examine connectivity between these two regions.

The tool region showed strong weighting of connections to visual regions, and accordingly, tools often have distinctive basic perceptual features ^30^. In line with tools often having distinctive perceptual features. Secondly, tools showed strong connections to the 4^th^ visual area, which is associated with color processing. Tools also showed strong connections to the third visual area, which is connected to posterior parietal regions that are associated with visuomotor transformations. Finally, the tool region’s strong connectivity with the posterior orbitofrontal complex (OFC) may be driven by top-down tool classification ^31^.

The HCP tool region was located in the cortex between the place and face regions. Although tool selectivity has been found before in this location ^32^ it is also present in other areas ^24^. From **Figure 1b**, it is apparent that even in the adults, there is some blurring between category boundaries, particularly between the tool and place regions. As the three classifiers were set up to each independently discriminate a single category selective region from all other voxels (including those that were selective for no category), these results cannot be used to quantify if pairs of categories can be distinguished from each other. To address this, we repeated the classification, but with a fitted discriminant analysis classifier that allowed for multiclass classification ^33^. Using multiclass classification meant a single classifier aimed to predict whether a voxel was face, place, tool or non-category selective. This confirmed that the three category-selective regions could be robustly discriminated from each other with even the smallest pairwise difference in d-prime, for tools vs. places, reliable in adults (*t*(13)=7.05, *p*<0.001) and infants (*t*(10)=2.40, *p*<0.05).

### Experiment 2

#### Category-Selective Regions Have Distinctive Signatures of Connectivity in Neonates

There are three limitations that Experiment 1 does not allow us to address. First, the sample size was small (N=11). Second, there was only one infant who was younger than three months. By three months, infants have had substantial visual experience, making it hard to distinguish brain representations that are innate from those that are learned. A larger sample of younger infants would allow us to identify the origin of early category-selective networks with a higher degree of between-sample generalizability. Third, the structural images from the infants in Experiment 1 were of insufficient quality to allow non-linear warping (normalisation), and so we used the fractional anisotropy images to register the adults and infants. There is the potential that this could mask differences in connectivity. In the adult group, we did have structural images, and we confirmed adult-to-adult classifications were similar if the structurals rather than FA images were used for registration. However, a more direct analysis would be possible if infant structural images were available for registration.

To address these three issues, in Experiment 2 we obtained diffusion weighted images from the second release of the Developing Human Connectome Project (dHCP) (N=400). The distribution of ages at birth and at scan is shown in **Figure 5**. As in Experiment 1, probabilistic tractography was used to identify connections from the ventral visual seed regions to target regions across the brain. The six linear discriminant classifiers trained on the adult data in Experiment 1 to identify face, tool and place selective regions in the two hemispheres were applied to the dHCP infant data. The classifiers were able to localize all three regions in neonates (Figures 6 and 7A, all p<0.001, faces: t(399)=70.89, places: t(399)=49.77, tools: t(399)=27.39). As for Experiment 1, we report results collapsed across hemispheres but each individual hemisphere was also significant. In contrast to our previous sample, faces and tools had a lower classification accuracy in the neonates than the adults in both hemispheres, respectively (faces: t(412)=2.32 p<0.05, places: t(412)=1.79 p>0.05, tools: t(412)=3.61, p<0.001) (**Figure 7**). This contrast with Experiment 1 could reflect less developed structural connectivity in neonates, or differences in scanners, sequences, or hemodynamics between the adult and neonate participants^34^.

**Figure 5.**
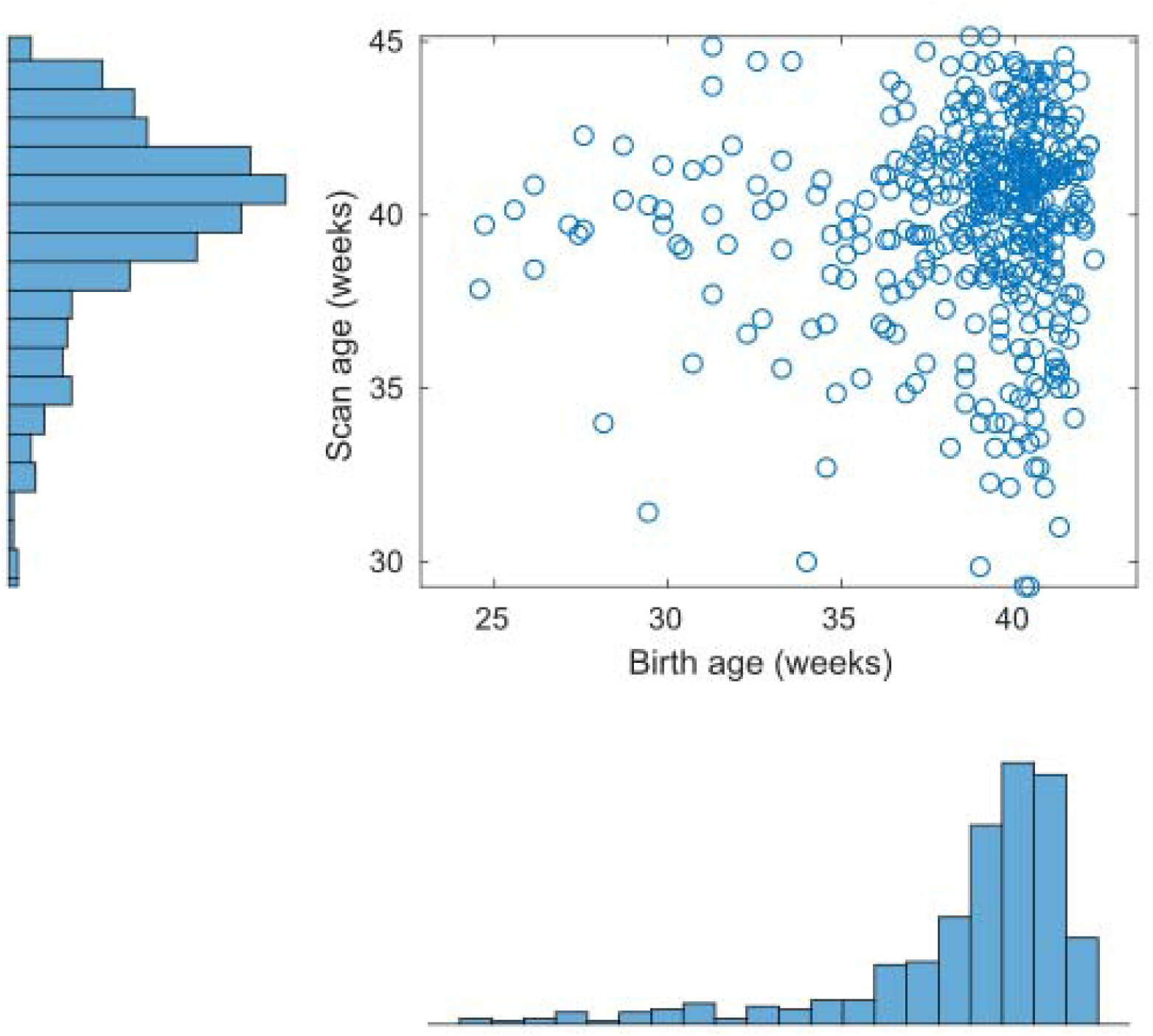
The postmenstrual age at birth and at scan for the 400 infants included in Experiment 2. The modal postmenstrual age of birth is 40 weeks.

**Figure 6.**
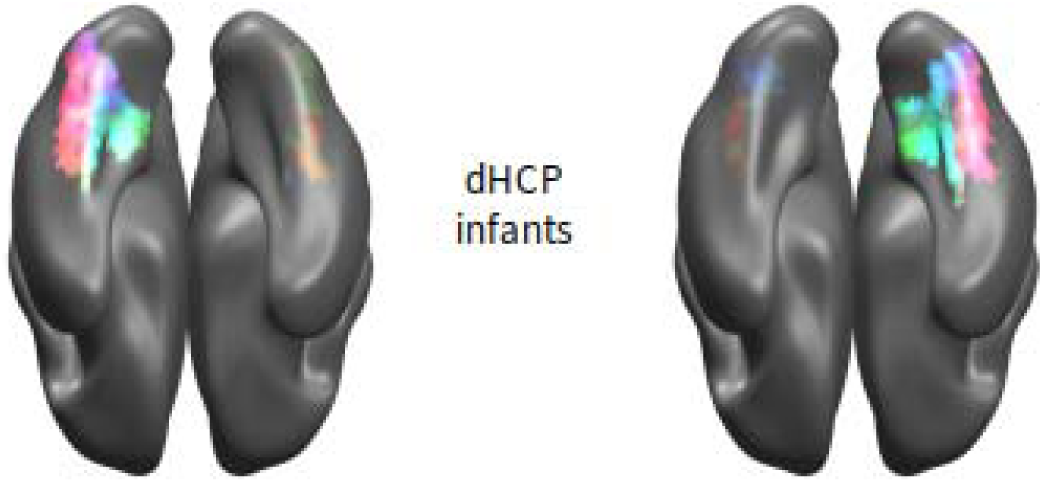
Distinctive signatures of structural connectivity were also present in neonates (N=400), as voxels could be identified as category-selective by a linear-discriminant classifier trained on adult connectivity and tested in infants. The projection is identical to Figure 2 (with red for faces, blue for tools, green for places).

**Figure 7.**
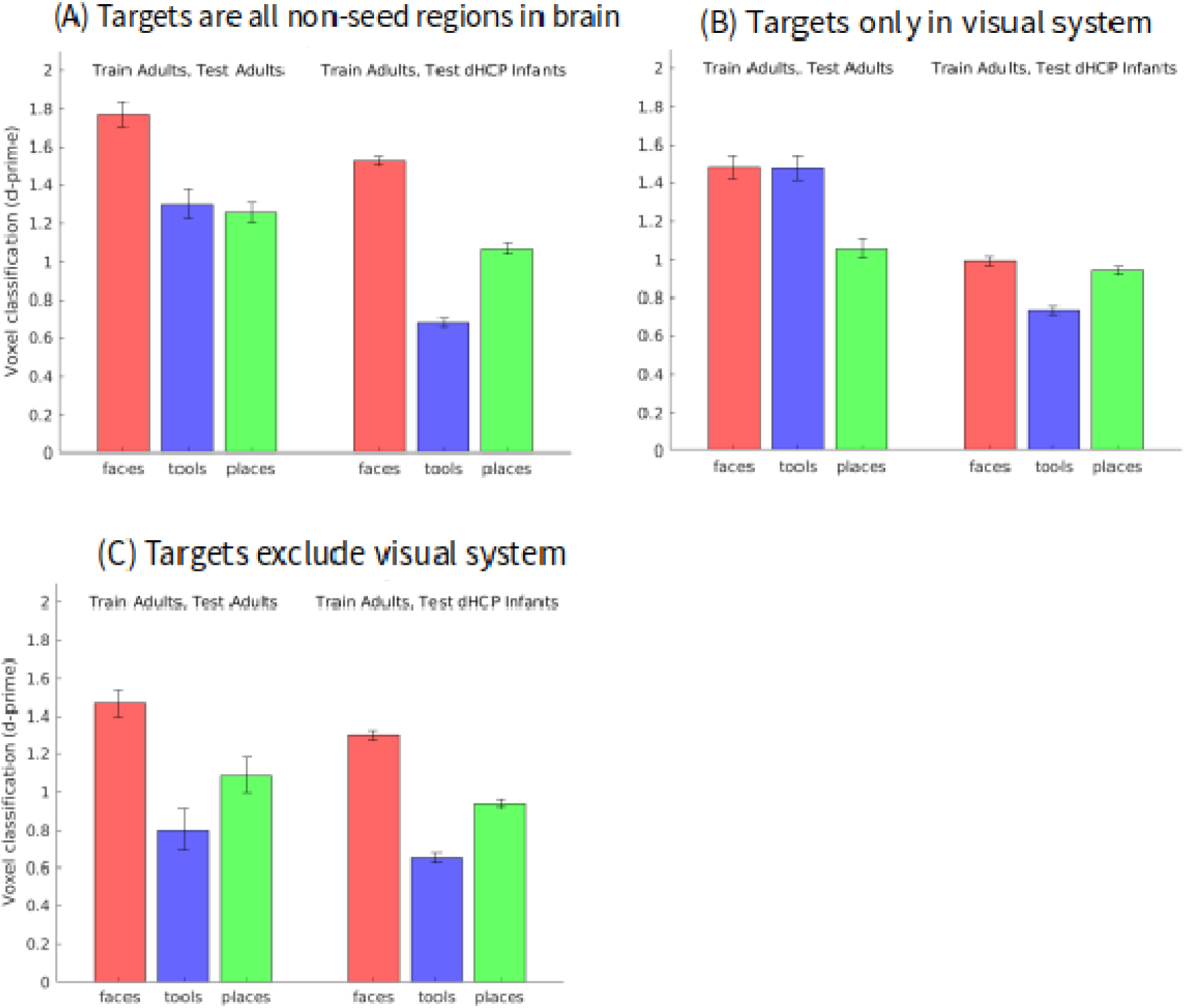
Voxel classification performance quantified using d-prime. a) Target regions for tractography were the 346 brain regions not included in the Glasser et al. (2016) ventral visual stream. All three regions were detected robustly in both the adults and neonates. In contrast with our previous sample, both the tool and place region were detected with a significantly less accuracy than adults. b) Target regions were restricted to only include those in the early visual system (V1, V2/V3 and MT+). All three category-selective regions could be localized in the adults and infants, demonstrating a proto-organization evident in early life c) Target regions were the inverse of b), containing all regions but early visual areas and ventral visual seed regions. The three category-selective regions were robustly localized in both the infants and the adults, demonstrating that long-range connectivity is present and part of the proto-organization in neonates.

#### Category selective connectivity to early visual regions vs. other brain regions

The results presented so far show for the first time using diffusion weighted imaging and tractography that category-selective regions have distinctive patterns of connectivity with the rest of the brain in young infants. This is consistent with the proto-organization of category selective regions, seen in infants with functional connectivity MRI by Kamps et al (2020). However, Kamps et al only studied connectivity between early visual regions and category selective regions, whereas in the results above we have used target regions throughout the brain. We therefore conducted two further analyses. In the first, we restricted the target regions to the early visual system - specifically, V1, V2/V3 and MT+ (networks 1, 2, and 5 in the Glasser atlas^26^). The three category selective regions in both hemispheres could be clearly identified from their connectivity pattern to early visual regions (**Figure 7B**) in adults (faces: t(13)=23.75 p<0.001, places: t(13)=22.32 p<0.001, tools: t(13)=21.61 p<0.001). for faces, places and tools, all p<0.001) and in infants (faces: t(399)=63.28 p<0.001, places: t(399)=81.69 p<0.001, tools: t(399)=56.50 p<0.001). These results concur with Kamp’s findings, extending them to tractography with diffusion weighted imaging.

In the third analysis, we selected the complement of target brain regions outside the early visual system (and of course, excluding the ventral seed regions). The three category selective regions in both hemispheres could again be clearly identified (**Figure 7C**) in adults (faces: t(13)=26.12 p<0.001, places: t(13)=22.27, p<0.001, tools: t(399)=17.17 P<0.001)) and infants (faces: t(399)=70.89 p<0.001, places: t(399)=49.77 p<0.001, t(399)=27.39 p<0.001). These results show that category-selective connectivity with distal brain regions is present early in infancy and suggest it forms part of an innate proto-organization.

#### Effect of Preterm Birth Suggests Connectivity of Category-Selective Regions is Innate

The previous result demonstrates the presence of a proto-organization containing both regions from the early visual system and long range connectivity to the rest of the brain. However, the effect of visual experience was not measured. The varying gestational ages in the dHCP data allow us to examine whether category-selective networks are experience dependent or innate, developing without visual experience. If the maturation of category-selective networks are experience dependent then infants who were born earlier will have greater visual experience and advanced development at the time of scan. In contrast, if category-selective networks are innate, they will not be dependent on visual experience, and preterm birth, as it places challenges on the infant, may delay development. To measure experience-dependent development, we calculated the time from birth until an infant’s scan. To measure innate development, we looked at the age of birth, while controlling for the time point at which the infant was scanned.

The relationship between d-prime and birth age, controlling for scan age was tested using partial correlation with Spearman (rank) correlation to address the non-normality of the age measures (see distributions in **Figure 5**). A small positive effect was found when considering connections that excluded the early visual system (r(400)=0.11, 0.03, 0.12; p<0.05, NS, 0.05 for faces, places and tools respectively) but not for connections only to the early visual system (r(400)=0.03, 0.03, 0.07, all NS). No relationship was found between experience-dependent development (i.e., the time between birth and scan) and d-prime, for either subset of connections (excluding early visual: r(400)=−0.05, −0.01, −0.08, all NS; visual only: r(400)=−0.03, −0.06, − 0.05, all NS). Taken together, this supports an innate, rather than experience driven, origin for the long-range connections for the face and tool networks.

### Experiment 3

#### Longitudinal Changes in Category-Selective Connectivity

In an independent dataset of N=45 neonates from the dHCP, two scans were acquired, separated by 6.8 +/- 2.4 weeks, with the postmenstrual age at the first scan of 34.5 +/- 1.7 weeks and at the second scan of 41.3 +/- 1.5 weeks. This provided the potential to examine longitudinal changes in the strength of category-selective patterns through this period. We found that when using this measure, there was evidence for the strengthening of connectivity with age for places (t(44)=1.78, 3.04, −1.23; p=NS, <0.005, NS for faces, places and tools).

## Discussion

The key results of the three experiments are summarised in Table 1. Category-selective connectivity was found to be both innate and shaped by experience. Connectivity of the face region was the most strongly innate as it was present in neonates, and was shaped by their age at birth, suggesting it develops *in utero* prior to visual experience of faces and is slowed by preterm birth. There was no evidence that the connectivity of the face region became more distinctive with experience in any of the experiments. Connectivity of place and tool regions was also found to be shaped innately, although only tools showed evidence of delay by preterm birth. However, places and tool connectivity did show evidence of developing with experience. Place-specific connectivity strengthened in the neonates, in the longitudinal study in Experiment 3, and tool-specific connectivity strengthened from 1-9 months, in Experiment 1. However, for both tools and places, we found no effect of the time between scan and birth (“experience age”) suggesting that either there was little development in this early time period in neonates, or that there are substantial individual differences that mask age effects in across-subjects analyses, but not within-subject longitudinal studies.

**Table 1.** Factors affecting category-selective connectivity.

|  | Exp. 1<br>1-9 months (N=11, 1 scan) |  | Exp. 2<br>Neonates (N=400, 1 scan) |  |  | Exp. 3<br>Neonates (N=45, 2 scans) |
| --- | --- | --- | --- | --- | --- | --- |
|  | Presence | Effect of age at scan | Presence | Effect of preterm birth | Effect of experience age | Longitudinal effect of age |
| Faces | *** | - | *** | * | - | - |
| Places | *** | - | *** | - | - | * |
| Tools | *** | * | *** | * | - | - |

The proto-organization previously found for the ventral stream and early visual regions was extended to include long-range connections with distal brain areas making up the adult unique signature of connectivity. The neonates, who had a modal postmenstrual age of 40 weeks, had adult-like connectivity patterns for distal brain regions excluding early visual regions. This evidence, in combination with the other results above, strongly suggests that the long-range connectivity of category-selective regions is not developed by a prolonged statistical learning of associations, but is instead largely innate, making up part of the proto-organization that had previously only been seen with early visual regions. Additionally, when looking at the effect of visual experience and birth age, only long-range connectivity was disrupted by preterm birth (for faces and tools), showing that within system network connectivity is less likely to be disrupted than between system network connectivity by early birth. The importance of long-range connections in semantics was recently demonstrated by multivariate decoding of white matter pathways in brain-injured patients with semantic deficits ^23^.

These results extend those from fMRI^35^ and functional connectivity^12^ in infants and structural connectivity in adults^14^ to reveal a proto-organization of category-selective regions in the ventral visual stream. The early maturity and specificity of the structural networks is further reflected in the connections identified by the classifiers. For example, many of the strongest connections for the place region are part of the place processing network—the hippocampus, entorhinal cortex and parahippocampal regions (see S1) ^2^ However, although we find consistent evidence of network maturity, there is substantial research that synaptic pruning continues into adolescence and that the microstructure associated with these networks continues to mature.

Although the results suggest that the face specific network matures early, even *in utero*, broadly mature structure does not imply that visual experience is not necessary to acquire a functional face representation. Both work in monkeys and with individuals who had cataracts as children demonstrate the need for experience to fully develop face processing capabilities, and infants under a month spend an estimated quarter of their waking hours with faces taking up the majority of their visual field 12^12,36^. There is, however, also some evidence of innate face processing. For example, Reid et al.^37^ demonstrated that fetuses will preferentially orient to face-like patterns in the third trimester. However, these results have generated controversy ^38^, and are not in line with multiple forms of evidence showing visual experience can shape the ventral stream.

In contrast to faces, place connectivity was not affected by age at birth, but did change longitudinally in neonates. There was also a lack of effect of “experience age” (time from birth to scan age) on connectivity, suggesting that place connectivity might not mature very early in life or that within-subject longitudinal analyses have more power to detect individual subject change. Regardless, it appears that experience may be important for the development of the place processing network. Young infants get substantial experience viewing scenes (and due to their long supine hours, perhaps particularly ceilings). The scene representations in the ventral visual stream are biased towards the periphery, and the retinal temporal hemifield, representing the periphery, develops before the nasal hemifield (representing more foveal representations) ^24,39^The place-selective network was found to contain structures like the entorhinal cortex, which is involved both in navigation and memory. It is likely that the complexity of creating multiple objects in space to form a coherent representation substantial enough to navigate requires a period of visual experience, and that the maturation of the visual system

Out of the three category-selective regions, the tool network was most consistently found to change with experience. Immature structure coupled with an infant’s tendency to see tools less than other visual categories could delay maturation. Young infants are perhaps less likely to see tools than faces and places. Infants younger than three months have primarily frontal views of faces, while older infants at 8-10 months have a mixture of body parts from nearby caregivers, some of which are acting on objects ^40,41^. These infants see a variety of objects at mealtimes, some of which are thought of as tools, and it has been suggested that the presence of these objects in the environment allow infants of the same age to look towards objects when they are named ^42^. These infants are able to sit up, crawl, and increasingly manipulate objects, although their tool-use behavior is not as sophisticated as that of toddlers.

Tools also are a less homogenous category, which will make category-level recognition more difficult, especially in comparison with faces, which have high similarity between exemplars. Compared with categories that are passively perceived, tool use also requires integration between sensory and motor representations, which might require more extensive experience with the environment. By 9 months, infants are able to differentiate between textures ^43^, and hold spoons correctly during self-feeding ^43^. Using head mounted eye trackers, researchers have determined that once infants learn to reach, they will often hold an object close to their face, which has been shown to be an ideal training stimulus for neural networks to recognize objects ^40^. In line with this evidence and the prolonged maturation of the tool network observed here, researchers have proposed a perception-action theory, where interactions between perception and motor experience gradually accrued over development explain the maturation of tool use ^44,45^. This theory is supported by behavioral evidence that experience with tools drives tool use behavior ^46^.

The machine learning technique developed in this work is, to our knowledge, used for the first time in infant neuroimaging. This method has the potential to create ‘growth charts’ or characterize the developmental trajectory of brain networks. In adults and children, connectivity has been shown to be predictive of many brain disorders, including neurodevelopmental disorders, and can be more predictive than the standard measures doctors will use to prescribe treatment ^47,48^. Early identification of an abnormal trajectory has the potential to facilitate early intervention when the infant brain is most plastic, which may be promising for potential infants, their families, and society as a whole.

A limitation of our results is the lack of functional localizers and diffusion data in the same participants, which would allow us to more accurately localize face place and tool regions in individual brains. Future research may be able to more closely identify the network maturation of category-selective regions with a comprehensive longitudinal study, where functional localizers are acquired in both awake infant and adult participants. However, one advantage to using the regions derived from the HCP is their generalizability across a large group of participants, something that would be challenging to do in a local sample. The HCP region definitions are also based on multiple types of data (structural, functional and diffusion data), which would also be challenging to acquire locally in large numbers in infants and adults. Nevertheless, these efforts would be worthwhile and could help answer many outstanding questions.

## Methods

### Experiment 1

#### Data Acquisition

For both the adult and infant participants, high-quality diffusion-weighted MRI data were acquired using a 3T Siemens Prisma Magnetron Scanner at the Centre for Functional and Metabolic Mapping of Western University. Using a 20-channel head coil, the Minnesota multiband sequence was used (128 directions, 2 mm isotropic, no gap between slices, b=1500 mm s^-2^, multiband acceleration 4, monopolar diffusion encoding gradients, time of acquisition: 9 min and 18 sec). Using monopolar diffusion encoding gradients creates larger eddy currents, which distort the magnetic field and cause image distortion. Using the solution developed by the HCP, two scans were acquired with opposite phase-encoding polarities (left-to-right and one right-to-left). Combining these images during analysis with FSL’s TOPUP calculates the susceptibility distortion, while EDDY corrects for eddy current-induced distortions and participant movement ^49^.

During the scan, younger infants were swaddled and wrapped in a Medvac pillow bag to help them remain still ^50^. Infants older than 6 months were not swaddled. All infants wore Mini Muffs adhesive sound protection (Natus, 7dB attenuation) and ear defenders (29 dB attenuation). Infants were scanned during natural sleep. Adult participants wore standard ear plugs and ear defenders and were requested to be as still as possible.

#### Participants

Diffusion-weighted MRI acquisitions were available from 11 sleeping infants as part of a larger infant imaging project with 51 participants. Infants were recruited either through public advertising or through clinical collaborators at the neonatal intensive care unit in London, Ontario. Diffusion MRI was acquired in 14 infants but three were subsequently excluded because of apparent brain injury. This left six healthy controls and four low-risk infants born preterm. One infant was scanned twice, but as the scans were two months apart, they were treated as separate participants in the analysis, making for a total of 11 infant datasets. Clinical information for the premature infants was obtained from medical records and a radiologist reviewed each scan for suspected brain injury. Infants were between 1 and 9 months old (corrected-age for infants born preterm, M=6.4 months, SD=3.2 months).

Diffusion-weighted MRI was also acquired from 16 adults at Western University. Participants were between 18 and 40 years old (M=22.75, SD=4.89). Author LC participated in the study and her data is included in the analysis. One participant was excluded because of an incidental finding, while another was excluded due to technical difficulties.

Approval for the study was provided by the Western University’s Health Sciences Research Ethics Board. All parents provided informed consent before infants were scanned. All adult participants also provided informed consent.

#### Preprocessing

The data was analysed with a pipeline built from the automatic analysis (*aa*) software^51^, FSL^52^, and custom Matlab (R2016a). *aa* divides the description of the analysis into a user script that describes what data should be analysed, the study specific settings, and a task-list. This user script then calls then aa engine, which runs the processing pipeline, ensuring that only the stages not already completed are run, and that when possible modules are executed in parallel. The task list describes which processing modules should be used to analyse the data. For this analysis, the modules identified the DICOM files and organized them based on header information (*aamod_autoidentifyseries_timtrio, aamod_get_dicom_diffusion*) and converted them to NIFTI format (*aamod_convert_diffusion_phaseencode_direction*). Then, *aamod_diffusion_extractnodif* identified the 10 volumes where b=0 in the diffusion data. The following six stages called components of the FSL diffusion processing pipeline. To combine the negative and positive phase encoding diffusion data into a single image and reduce distortion, *aamod_diffusion_topup* (TOPUP). *aamod_bet_diffusion* then removed non-brain tissue in the b=0 image (BET). In order to correct for any residual distortions due to eddy currents or head motion, *aamod_diffusion_eddy* (EDDY). *aamod_diffusion _dtifit* was then run to model diffusion tensors at the voxel level (DTIFIT).

In order to obtain mappings between individual brains to standard (MNI) space for the infants and adults, the normalization procedure for FSL’s tract-based spatial-statistics (TBSS)^53^ was run. This normalizes the fractional anisotropy (FA) image to a mean FA tract skeleton, using non-linear registration. Normalizing the FA image resulted in a good registration for both the infant and adult data. Conventional normalization using a structural (T1 or T2) image to a template was not possible for the infants, as a number of the structural images were of poor quality due to participant motion. To ensure that normalizing with FA was not introducing an artifact into our results, additional analyses (not shown) in the adults where good structural images were also available, confirmed that very similar results were obtained if normalization was performed using the structural rather than diffusion images.

#### Human Connectome Project

In order to identify seed and target regions for the diffusion analysis, the parcellation from the HCP was used. The HCP parcellation segments the brain into 180 distinct regions in each hemisphere, based on structural, functional and diffusion data. To identify the seed regions, the HCP definition of the ventral stream visual cortex (region 4, supplementary neuroanatomical results) was used to identify the 14 regions that make up the ventral visual stream ^26^. The individual voxels that were part of the 14 regions in the ventral visual stream were used as seeds and were excluded from tractography targets. The other 346 regions from the parcellation served as the target regions in the analysis. These seed and target regions were projected from the cortical surface into volumetric MNI space. The normalization parameters from TBSS were then used to project these regions from MNI space into each subject’s individual diffusion data space for tractography.

To select the regions in the ventral visual stream which were most responsive to faces, places, and tools, the functional MRI localizers from the HCP project were used. The category-average contrasts were used to select regions. These regions were the fusiform complex, the ventromedial visual area 2, and ventromedial visual area 3, respectively. Regions can be seen in **Figure 1A**. These regions were used as the category-selective regions in the subsequent classification analysis.

#### Tractography and Classification

Using the data from aa’s *aamod_diffusion_bedpostx* module (BEDPOSTX), probabilistic tractography was performed in the infants and adults using FSL’s PROBTRACKX, using 5,000 streamlines per seed voxel in the individual subject’s ventral visual stream. The output of PROBTRACKX was then transformed to MNI space. These results were then summarized into a connectivity matrix that contained, for each voxel in the MNI ventral visual stream seed region the number of streamlines that terminated in each of the 346 target ROIs.

Three linear discriminant classifiers were then trained to identify the fusiform complex, the ventromedial visual area 2, and ventromedial visual area 3 in adults, based on connectivity with the 346 target regions. For the adults, leave-one-subject out cross validation was used to test whether selectivity could be predicted from connectivity. For the infants, three classifiers trained in a similar way on the full adult dataset were then tested on the infant diffusion data. D-primes were calculated to evaluate the accuracy of the classifiers in both infants and adults. To test for a relationship between age and connectivity in the infants, Pearson correlations were calculated between age and d-prime scores for each category.

For the multiclass classification, a fitted discriminant analysis classifier (www.mathworks.com/help/stats/fitcdiscr.html) was used to identify the fusiform complex, the ventromedial visual area 2, and ventromedial visual area 3, as well as the non-category selective voxels in the HCP ventral stream visual cortex (region 4, supplementary neuroanatomical results) in adults. This was done based on voxel-wise connectivity with the 346 target regions. Leave one-out cross validation was used to test classification accuracy, and d-primes were calculated for each participant. For the infant version of this analysis, the fitted discriminant analysis classifier was trained on the entire adult dataset and tested on the infant data. D-primes were used to calculate classification accuracy.

#### Statistics

For the single class analysis, the following statistics were calculated. Six one-sample, two-tailed t-tests were conducted (one for each category, in both the infants (N=11) and adults (N=14)) to test if the d-primes were reliably different than zero. A repeated-measures ANOVA examined main effects of and interactions between age group (infants (N=11) vs adults (N=14)), hemisphere, and category. Three two-tailed, repeated-measures t-tests were conducted to compare adult d-primes to infant d-primes; one test was conducted per category. Three Pearson’s correlation coefficients were calculated to test if there was a relationship between age and d-prime in the infants (N=11); one was conducted per category. One paired-samples, two-tailed t-test was conducted to compare the hit rate between the place and tool region in the adults (N=14). For the multi-class analysis, tool d-primes were compared to place d-primes in a paired-samples, two-tailed t-test. This was done for both the infants (N=11) and adults (N=14).

### Experiment 2

#### Dataset: Developing Human Connectome Project (dHCP)

The dHCP is a unique project with multiple imaging modalities (structural, functional and diffusion images) combined with behavioural and genetic data. Our analysis is based on the second data release with 558 neonates, which yielded 490 sessions of usable diffusion imaging data from 445 unique participants - 400 with a single session and 45 with two sessions. This release included both term and preterm infants with gestational ages ranging from 24-45 weeks. In experiment 2, we report analyses from the 400 infants with a single scanning session. Infants were scanned while sleeping (without sedation) at the Evelina Newborn Imaging Centre, St Thomas’ Hospital, London, UK. Ethical approval for the study was provided by the UK Health Research Authority.

#### Diffusion

Neonates were imaged on a 3T Phillips Achieva scanner using a customized 32-channel neonatal head coil ^54^. Diffusion weighted images were acquired using an optimized multi-shell HARDI sequence which varied the number of directions on four shells (b0, b400, b1000, b2600) designed to optimize the white and grey matter contrast in neonates. All four phase encoding directions ^55,56^ were used and the image resolution was 1.5mm isotropic. The scan lasted approximately 20 minutes. If the neonate woke up during the scan and subsequently settled, the scan was restarted to overlap with the previous acquisition. More details regarding the diffusion acquisition can be found in Bastiani et al. 2019 ^57^.

#### dHCP diffusion preprocessing

Diffusion data obtained from the dHCP was preprocessed using the pipeline from Bastiani et al. (2019)^57^. Susceptibility induced distortions were corrected using estimation from the four PE directions acquired. FSL’s EDDY was used to correct for additional distortions including those caused by eddy currents, and motion. Outlier slices were detected and replaced and slice-to-slice motion correction was performed. A super resolution algorithm was applied to resample the thicker slice spacing (3 mm) onto the isotropic grid (1.5 mm).

#### Diffusion Quality Check

As part of the dHCP release, all diffusion data underwent a quality check. Metrics were derived from the EDDY QC tool. The SNR values as well as three contrast to noise ratios, one for each of three b values (b400, b1000, b2600) were z-scored against all participants, and those who had a z-score of less than-2.0 were excluded by the dHCP leaving 490/558 participants.

#### Transformation of HCP ROIs to dHCP individuals

As for Experiment 1, the seed and target ROIs for tractography were generated using the parcellation of Glasser et al (2016). Tractography was performed in the native space of each infant using FSL’s probtracx2. To facilitate this, the seed and target ROIs were transformed from the adult space to the native infant space. The dHCP release included the transformation from the individual diffusion image space to Bozek et al’s ^58^ 40 week template in volumetric space (https://gin.g-node.org/BioMedIA/dhcp-volumetric-atlas-groupwise), which could was applied using FSL’s applywarp with nearest neighbour interpolation. To calculate the transformation from this dHCP 40 week T1-weighted template to the skull-stripped adult T1-weighted template MNI-152 space, we used ANTS, as it has previously proven effective in pediatric studies ^59–61^. In assessing registration, we initially found there were two problems. One was that the scalp in the infant brain is thinner and closer to the cortex (see Supplementary **Figure 2a**) and the adult cortex would register to this. We therefore removed the scalp from both the infant images by taking the background tissue (type 4) as provided with the atlas, thresholding it (threshold 1000 determined by eye), inverting this mask, and applying it to the infant T1. A second issue was that relative to the cortex the cerebellum is smaller in infants. In initial attempts, this led to the inferior visual cortex being dragged down during the non-linear warping process. We therefore masked out the cerebellum in the adult and infant templates. For the adults, we downloaded the AAL3 template ^62^ from https://www.oxcns.org/aal3.html and created a mask of the cerebellar regions 95-120, inverted it, and applied the mask to the T1 template. We masked the infant cerebellum by using tissue type 6 from the 40-week atlas, thresholding by 300, inverting, and masking the 40-week T1-weighted image. The consequent ANTS registration is shown in Supplementary **Figure 3**.

#### Tractography

Tractography analyses were conducted in the AWS cloud using a cluster made using parallelcluster (https://docs.aws.amazon.com/parallelcluster/), with SLURM as a queue manager. GPU compute instances were used for accelerated CUDA 10.0 versions of bedpostx and probtrackx (CITE https://journals.plos.org/plosone/article?id=10.1371/journal.pone.0130915) downloaded from https://users.fmrib.ox.ac.uk/~moisesf/Probtrackx_GPU/Installation.html. Bedpostx was run in the same way as reported for the dHCP pipeline ^57^ https://git.fmrib.ox.ac.uk/matteob/dHCP_neo_dMRI_pipeline_release/-/blob/master/dHCP_neo_dMRI_runBPX.sh) Probtrackx was run using seeds and targets defined in the same way as for Experiment 1. During the registration to native space, some ROIs would fall outside the field of view of the acquisition. These target ROIs were not included as targets for that baby. Probtrackx2 was then run using 5000 streamlines, with the switches --onewaycondition, --opd and --os2t and --rseed=1234.

#### Transformation back to MNI space

For each target, an image was produced that described for each seed voxel, what number of streamlines ended up in that target. These images were transformed back from the native space to the 40 week template using FSL’s applywarp, and then from the 40 week template space to the MNI space using ANTS. The classification analyses could then be performed on the seed-to-target matrix in MNI space, which was common to all of the infant and adult participants.

#### Connectivity signature analysis

As for experiment 1, in the adults, classification was performed with leave-one-subject-out cross validation. Each hemisphere for each of the three categories (faces, tools and houses) was taken in turn. In each cross-validation fold, a linear-discriminant classifier was trained to discriminate seed voxels that were or were not selective for the category based on the connectivity of that seed to the targets. It was then tested on the left out subject and the signal detection measure of d-prime used to summarise detection performance.

To test whether proto-organization in connectivity was present for just the visual cortex, or also other brain regions, we conducted two additional classification runs. In one the set of targets excluded the early visual regions categorized as V1, V2/V3, ventral stream and MT+ (i.e., excluded networks 1, 2, 4 and 5 from the 22 networks labelled by Glasser et al, 2016). To assess the classification with only visual regions, we included only V1, V2/V3, the dorsal and ventral streams and MT+ (regions 1-5 from the 22 networks).

#### Gestational and scan age effects

To test whether there were effects of age-at-birth or age at scan, the d-prime measurements were calculated as described above for and excluding early visual regions. Then partial Spearman (rank) correlations were calculated for birth and scan age and d-primes measurements based on data that excluded or included targets in the early visual system. Spearman (rank) correlations were used to address the non-normality of the age measures.

### Experiment 3

For Experiment 3, we selected an independent subset of the dHCP dataset described in experiment 2, of the 45 neonates with two scanning sessions. The analysis methods were identical, except that the only outcome measure of interest was the difference between the two scans in the d-prime classification performance for each visual category collapsed across hemispheres. This was calculated using a paired t-test, although qualitatively identical results are obtained using a non-parametric Wilcoxon rank sum test.

## Data Availability

The data from the HCP are available from https://www.humanconnectome.org/study/hcp-young-adult/data-releases.. Raw fMRI data has not been made publicly available because it is sensitive medical data, and the ethics that the data was collected under did not specify that we would store it in a public repository. This is especially relevant because our study contains data for infant participants, whose parents consented to their participation in our study. Instead, we will make the matrices that the classification was performed on publicly available.

## Code Availability

Custom code used in the study will be made available in a Github repository. All FSL and *aa* software is already publicly available.

## Acknowledgements

From the dHCP: “Data were provided by the developing Human Connectome Project, KCL-Imperial-Oxford Consortium funded by the European Research Council under the European Union Seventh Framework Programme (FP/2007-2013) / ERC Grant Agreement no. [319456]. We are grateful to the families who generously supported this trial.”

## Supplementary figures

**Table S1.**
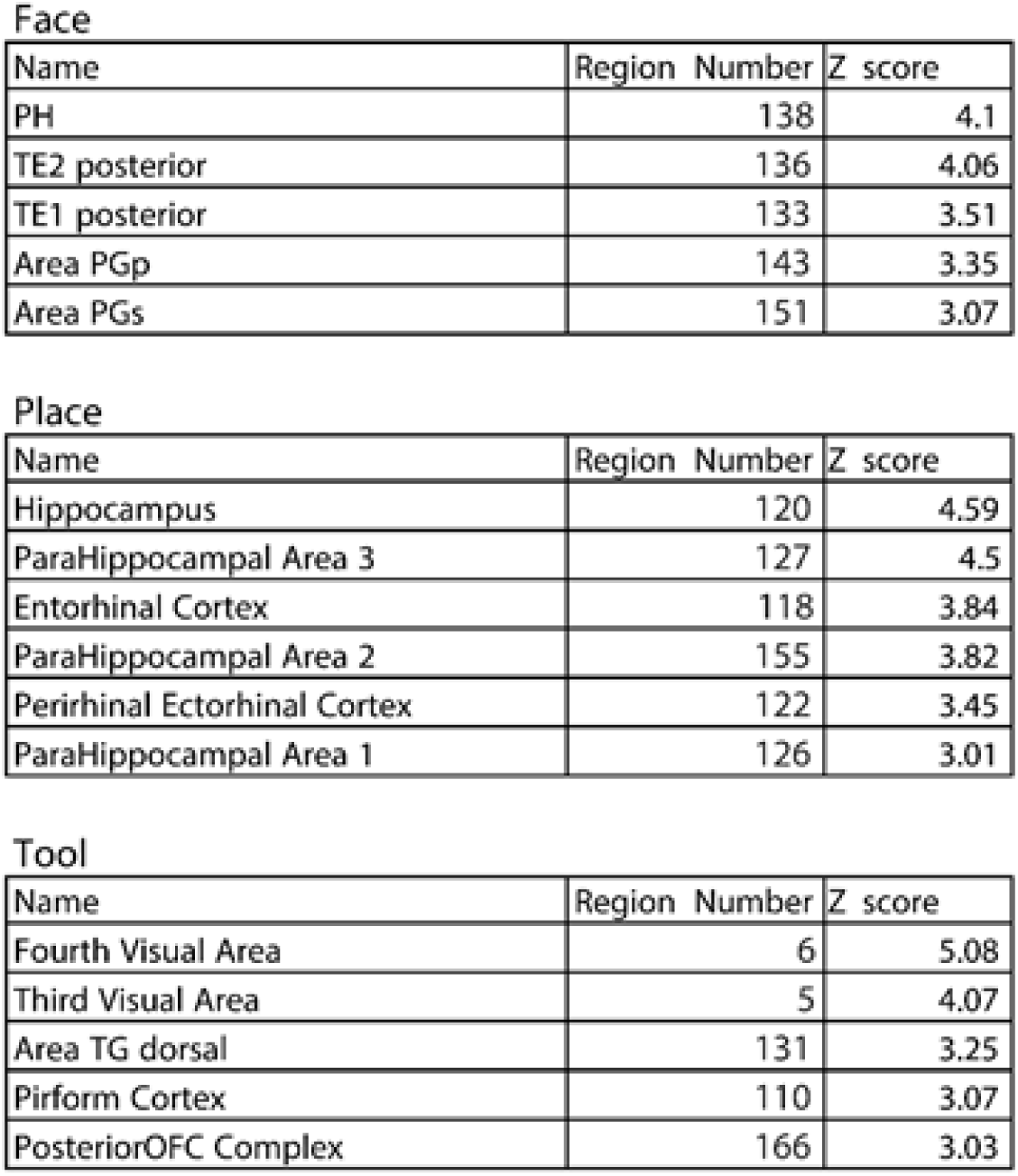
In order to determine which regions carried the most signal for classification, the mean difference in connectivity between category-selective and non-category selective seed voxels was calculated for all target regions. It was standardized across target regions to yield a z-score. This table shows regions with a z-score that is greater than 3.

Face
| Name | Region Number | Z score |
| --- | --- | --- |
| PH | 138 | 4.1 |
| TE2 posterior | 136 | 4.06 |
| TE1 posterior | 133 | 3.51 |
| Area PGp | 143 | 3.35 |
| Area PGs | 151 | 3.07 |

Place
| Name | Region Number | Z score |
| --- | --- | --- |
| Hippocampus | 120 | 4.59 |
| ParaHippocampal Area 3 | 127 | 4.5 |
| Entorhinal Cortex | 118 | 3.84 |
| ParaHippocampal Area 2 | 155 | 3.82 |
| Perirhinal Ectorhinal Cortex | 122 | 3.45 |
| ParaHippocampal Area 1 | 126 | 3.01 |

Table S1. In order to determine which regions carried the most signal for classification, the mean difference in connectivity between category-selective and non-category selective seed voxels was calculated for all target regions.
| Name | Region Number | Z score |
| --- | --- | --- |
| Fourth Visual Area | 6 | 5.08 |
| Third Visual Area | 5 | 4.07 |
| Area TG dorsal | 131 | 3.25 |
| Piriform Cortex | 110 | 3.07 |
| PosteriorOFC Complex | 166 | 3.03 |

**Supplementary Figure 1.**
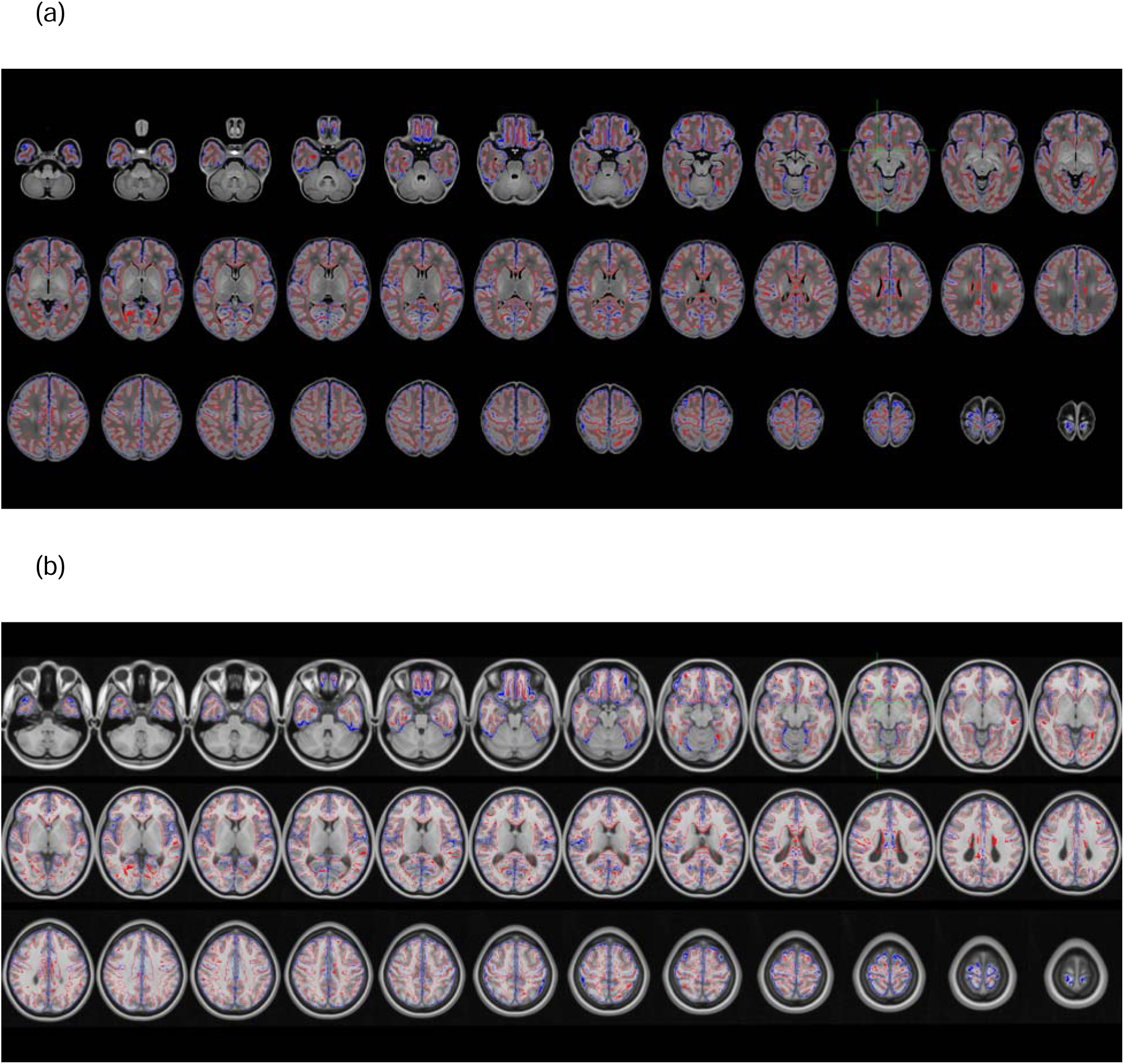
Final registration of infant and transformed-adult spaces. (a) The infant 40 week T1-weighted template (greyscale images) with the pial boundary (blue lines) and the grey-white boundary (red lines). (b) The adult T1-weighted template warped into the 40 week template space using ANTS (greyscale images), shown with the pial and grey-white boundaries from the infants (blue and red lines, respectively). Registration was generally of good quality.

**Supplementary Figure 2.**
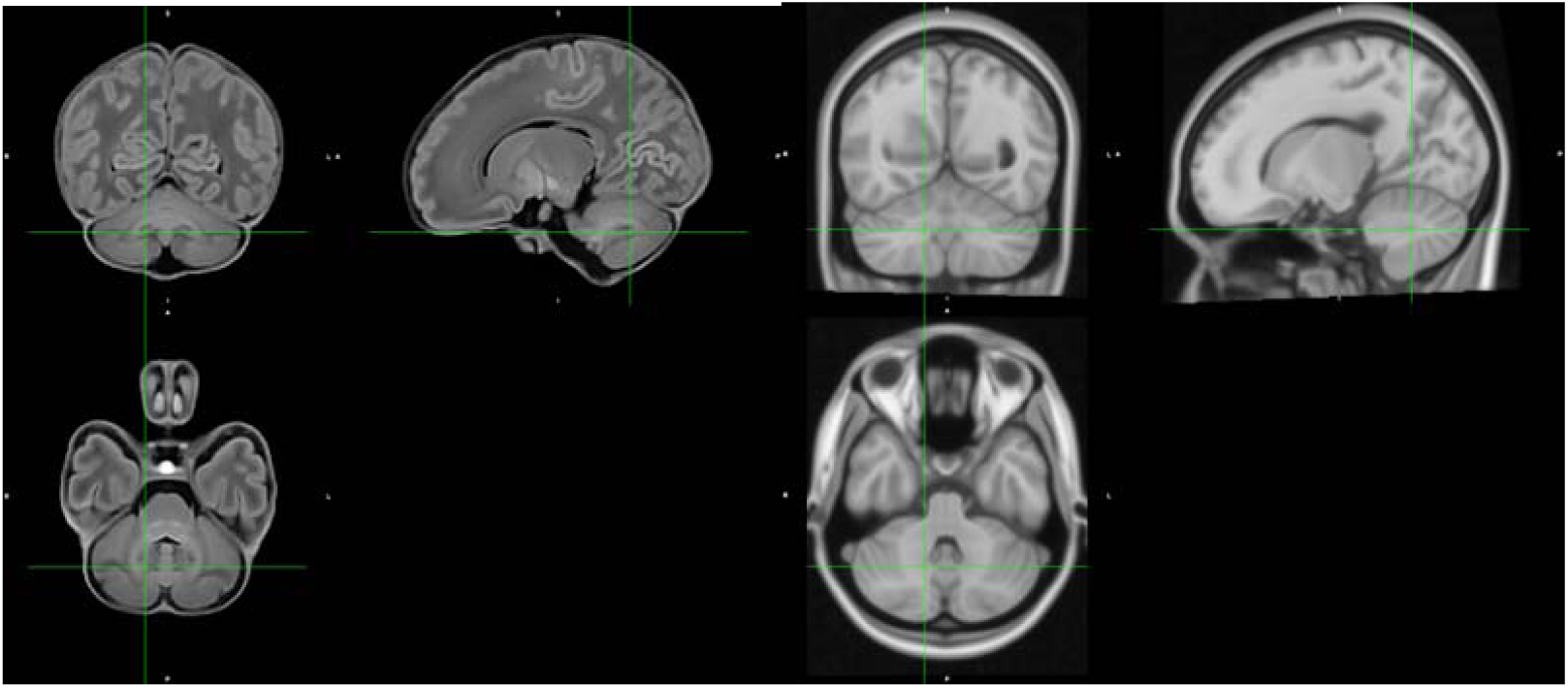
Relative to the size of cortex, the cerebellum is smaller in infants (left) than in adults (right)

## Notes

### Competing Interest Statement

The authors have declared no competing interest.

